# Topology Adaptive Graph Convolution Network With Heterogeneous Entities For Predicting Adverse Events from Drug-Drug-Interactions

**DOI:** 10.1101/2022.05.16.491112

**Authors:** Srijani Bagchi, Anasua Sarkar, Ujjwal Maulik

## Abstract

In times like this, it is imperative to be cautious about the effects of drugs or vaccination doses on patients who are already suffering from other serious diseases. It’s not only the virus which can affect the body metabolisms, drugs to encounter the virus may also end up having unwanted negative effects. Therapeutic activities of drugs are often influenced by co-administration of drugs that may cause inevitable drug-drug interactions and inadvertent side effects. Therefore, prediction and identification of DDIs are extremely vital for patient safety and treatment modalities. In this context, we intend to develop a computational method based on functional similarity of drugs. Our objective has been enhanced by the usage of knowledge graphs which allows capturing underlying information embeddings in a biological network with heterogenous entities. On providing this knowledge graph as the input to a Topology Adaptive Graph Convolution Network which performs topologically-aware flexible convolutions, we achieve improvements on priorly proposed GCN models that has been shown as comparison.

## 1. Introduction

Predicting drug-drug-adverse events is a complex task because it requires figuring out the underlying mechanism of action of two drug interactions. The interaction of drugs depends on many factors like genes/targets, diseases, and dosages of drugs. There exist numerous many-to many relevant associations among entities like a drug, target/gene, adverse event, etc. In recent years, representation learning plays a significant role in many data science domains, such as natural language processing, computer vision, etc. They convert the structure of the input data as an element in a low dimensional vector space, however in some domains, forceful regularization of data causes the model to lose its ability to learn the structure of input signal. Hence, all the data is assembled by constructing a knowledge graph on top of the existing designed relational knowledgebase. The acquired information from the knowledge base and knowledge graph are integrated into a framework.

Convolutional neural networks have been effective in identifying patterns in data. The learning ability of these models are owed to the kernels that learn local, translation-invariant functions over the input signal. When stacked layer upon layer, CNNs can extract low-level features and construct high-level representations of the input data. However, CNNs are suited to domains where data is structured and regular.

However, many domains exhibit an irregular structure that does not allow conventional application on convolution for feature representation purposes. Irregular data can be forced to be made regular, but the model loses the ability to learn the structure of the input signal ^[1]^. Such irregular data structures are commonly represented in the form of graphs. Various types of biological data are commonly represented as graphs. These include relationship between genes, association between diseases, drugs and so on. In such cases, graphs are not guaranteed to have local or regular structures, and hence, conventional convolution and pooling operators do not work with graphs. Difficulties arise during representation learning on graphs since conventionally used operators cannot be applied directly. Initially, domain specific properties were developed for developing convolution ^[2]^, but later, the focus shifted on more generic techniques ^[3]^.

## 2. Literature Survey

In recent times, large knowledge graph (KG) plays a crucial role in various applications by utilizing valuable information ^[4]^. The emerging importance of KG and improvement in many ma-chine learning techniques is an important factor in choosing this area. In the biomedical domain, a specific entity based small graph is often designed to accomplish certain tasks like gene-drug, drug-drug predictions, etc. Therefore, the design of a large-scale biomedical knowledge graph can be effective and necessary in solving many realistic cases.

The KG can be described as the acquisition of relational facts from relevant sources. KGs add values to many question-answering ^[5]^ and search applications that require understanding semantics between real-world data. Properly curated prior knowledge is represented in the form of entities (nodes) and relationship (edges) in the KG. It allows machines to understand the semantics among entities and figure out the underlying concepts. There have been variations in the context in which KGs have been used. In the spatial domain, the core mechanism of graph convolution and its variants rely on neighbourhood aggregation. This process aggregate neighbours to compute hidden node representation of a graph. The aggregation process relies on message passing techniques where node features are transformed and passed from the neighbouring nodes and are subsequently aggregated. State-of-the-art architectures such as MoNet, Patchy-SAN, GCN, GraphSAGE, GAT or GIN incorporates these techniques.

Several limitations are encountered by researchers for the construction and completion of the KG. For starting KG construction, the basic requirements are concept knowledge base and substantial knowledge source. However, it requires expert knowledge and time to gather exhaustive domain knowledge in the biomedical domain, which itself is a challenge. The way to overcome such challenges is by designing a framework that has the facility to access information from one source. Such a source congregates information by integrating multiple heterogeneous sources. Thus, allowing researchers and domain experts to investigate information best suited as per their requirement.

Over the years, most of previous works are proposed to predict the potential DDIs by either integrating multiple data sources or combining the popular embedding methods. Different from drug similarity obtained from multiple sources ^[6]^, a deep learning framework named Deep-DDI ^[7]^ is proposed to use molecular structures of drug as inputs, and to predict additional DDI types. In the same line of work, ^[8]^ integrates several graph embedding methods for DDI task, and with the assistance of knowledge graph, ^[9]^ models DDI as link prediction. With comparison to the classic and graph embedding methods, our proposed framework is able to automatically extract drug features from the data, and requires neither chemical structure nor specialized drug representation.

Recently, there has been an increasing interest in applying graph neural networks for DDI prediction. To effectively aggregate the feature vectors of its neighbours, different aggregation strategies lead to different variants of GNNs. *Decagon* ^[10]^ applies a relational GNN for modelling polypharmacy side effects. To extract DDIs from text, ^[11]^ utilizes a graph convolutional network (GCN) to encode the molecular structures. However, we have referred to the original and basic algorithm which entails generalization of the convolution operation for non-Euclidian space. The modifications are performed on the filtering operations and other network model details as will be discussed later.

Machine Learning Models are data-driven by nature. They are specified by models that relate one or more independent variables to other dependent variables. These models provide prediction by utilizing known properties or patterns learned from the training data. The models are also involved in the discovery of unknown or new patterns learned within the data. Data driven learning problems are formulated as minimization of some loss function based on the available training set. The loss function provides the difference between the actual and predicted values, which is utilized to tune the model further to improve the model’s accuracy. Moreover, the core objective of a learner is to generalize from its experience. These models usually learn the representations from large amounts of available biomedical data and have performed predictions from the data’s acquired characteristics.

## 3. Problem Definition

The main objective of this work is to propose a classifier-based learning model to predict adverse events from combined drug interactions. The over-all approach of binary classification task of adverse events comprises of three essential steps: (i) acquiring data, (ii) preprocessing data, and finally, (iii) classification ofdata.

The problem of adverse event prediction is modelled as a binary classification task. Given two drugs *d*_*p*_ and *d*_*q*_, the target-label is a binary classifier. For *Y (dp,dq)* is 0 indicates drug interaction does not happen, whereas *Y (dp,dq)* is 1 indicates the drugs *d*_*p*_ and *d*_*q*_ interact to produce a side effect, which might be adverse or beneficial.

## 4. General Frameworks

The basic concepts whose applications have been implemented to give a direction to this research are described in this section:

### 4.1. Drug Interactions

The science of drugs is known as pharmacology. In living systems, it deals with molecular inter-actions that produce a biological response. When simultaneously, two or more drugs are given for treating a disease, then the therapeutic effect of one drug action may get modified by another drug. Such type of effect of the drug occurs due to drug-drug interaction (DDI). The drug-drug interaction effect can either result in positive catabolism or in a harmful adverse reaction. Drug-drug interactions can occur outside (in vitro) and inside (in vivo) the body.

Predicting adverse event from co-prescribed drugs are of high significance before releasing the drug into the pharmaceutical market. An application of drug-drug-adverse event prediction is analysed using integrated data sources. A network of protein-drug is designed to learn embedding for each drug entity. Classifying drug-drug interactions mechanistically gives major insights into how to predict, detect and avoid them:

- **Pharmaceutical interactions:** Such interaction can occur due to the physical or chemical in-compatibility of a drug with the external syringe infusion. For example, applying phenytoin in dextrose can result in precipitation; loss of potency can occur if carbenicillin and Gentamicin are infused together.
- **Pharmacokinetic interactions:** In Greek, “kinesis” means movement, i.e., the action of the body on alteration of drugs. Such alteration includes absorption, distribution, biotransformation, and elimination of drugs. For example, within 30-60 minutes paracetamol gets absorbed orally; renal tubular secretion of penicillin/cephalosporins is decreased byprobenecid.
- **Pharmacodynamic interactions:** In Greek, “dynamics” means power, i.e., the action of the drug on living system/body. Results in the biochemical and physiological action of chemical molecules/drugs at systems like an organ at subcellular/macromolecular levels. Such type of interaction may result in additive, antagonistic or synergistic effects. For example, parkinsonism has a beneficial effect by combining levodopa and carbidopa; the interaction of aminoglycosides and amphotericin may result in a harmful effect by enhancing nephrotoxicity.
- **Absorption:** By affecting the dissolution of the drug in the stomach, by influencing gastric emptying or intestinal blood flow or by inhibition of active transport processes, certain interactions can increase or decrease the rate of absorption.
- **Distribution:** Protein binding, tissue binding
- **Metabolism:** Hepatic/Non-hepatic
- **Excretion:** Renal/Non-renal

### 4.2. Knowledge Graphs

A knowledge graph is a model of storing domain knowledge data like entities (concepts, events, situation, object) interlinked with each other as the relationship between facts. It can be designed based on labelled property graphs (LPG) or resource description framework (RDF) formats. It is mainly built on the top of the existing database.

The top-down approach of knowledge graph construction emphasizes on domain knowledge ontology schema. In this study, the top-down approach to design a biomedical knowledge graph is employed. According to that method, the designed knowledge graph schema is defined first, then populates the graph with its instances. For the construction of KG into a significant network format, the property graph model is exploited. The property graph model means the data in the graph database are organized in the form of nodes, edges, and properties. The nodes are labelled with broad category names based on the underlying concepts/facts. The edges are represented with directions based on the semantic associated among nodes.

A systematic way to interconnect key concepts is necessary. In the biomedical domain, it plays an important role in many problem solutions such as interaction prediction, missing link prediction, and inferencing algorithms. The construction of the biomedical graph is from the assembled data is essentially the essence of knowledge graphs. The graph nodes are represented by concepts (entities) are of three types. The biomedical entities can be drugs, diseases and genes. The semantic relation between those concepts (relations) is also labelled.

The schematic representation of the integrated knowledge graph is shown in Figure 1. The oval boxes represent disease, drug or gene entities. The rectangular boxes highlight the properties or features of each type of entity. The line connecting the two nodes show the type of relationship between them. The relationships are named as *interacts with, targets, treats*. The relationship type can be either one-directional or two-directional based on underlying semantic between the nodes/concepts.

**Fig 1:**
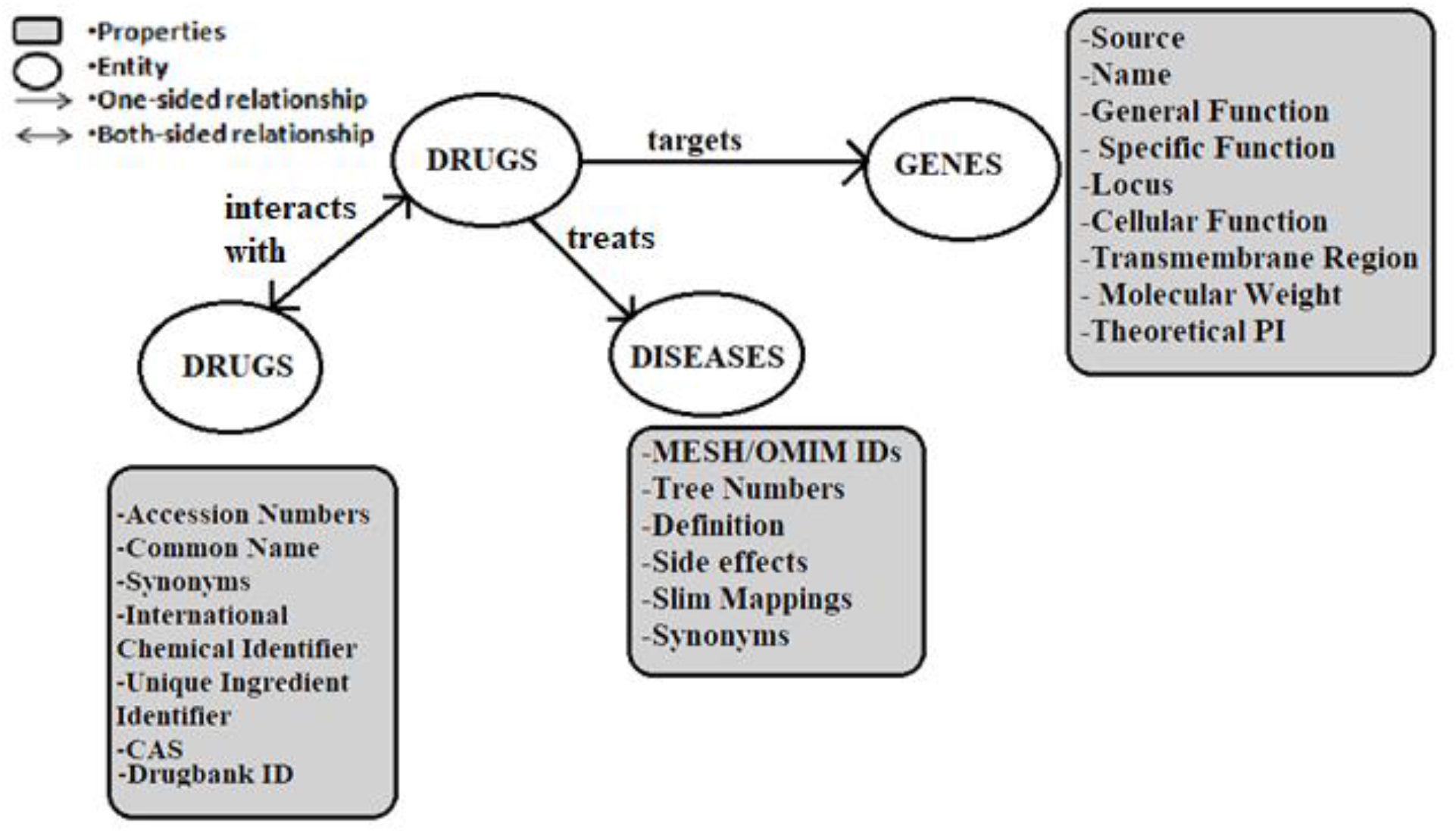
Schematic representation of the integrated Knowledge graph model.

### 4.3. Topology Adaptive Graph Convolution Network

In order to learn non-linear representations embedded in the knowledge graph-structured data, we have implemented topology adaptive graph convolutional network (TAGCN), a unified convolutional neural network proposed by Du et. al ^[12]^. It slides a set of fixed-size learnable filters on the graph simultaneously, and the output is the weighted sum of these filters’ outputs, which extract both vertex features and strength of correlation between vertices. Each filter is adaptive to the topology of the local region on the graph where it is applied. TAGCN unifies filtering in both the spectrum and vertex domains; and applies to both directed and undirected graphs.

In graph signal processing, graph shifting is crucial to incorporate adaptive graph filtering. Sandryhaila and Moura showed in ^[13]^ that traditional discrete signal processing can be extended to biological networks by context-appropriate interpretations of shift-invariance and invertibility properties of graph filters.

The proposed TAGCN scans the graph before performing the convolution, and based on the topology of the graph, decides the weights and topologies of the filters. Though the convolution for the vertex domain is consistent with convolution in traditional CNNs, the fixed square filters in traditional CNNs for grid-structured input data volumes are replaced with K-localized filter for graph convolution to extract local features on a set of size 1 up to size K receptive fields. This is because on analysing the mechanisms of graph convolutional layers, it can be realized that when the convolutional layers go deeper under certain conditions with only a size k filter, the output of the last convolutional layer is the projection of the output of the first convolutional layer along the eigenvector corresponding to the eigenvalue of the graph adjacency matrix with largest amplitude. By avoiding this linear approximation, we also get rid of information loss classification accuracy degradation. So, using a set of size 1 to size K filters increases representation capability, which leads to improved classification accuracy. It manages to achieve these in lower computational complexity than the basic GCN algorithm or its recent modifications. This can be attributed to the fact that TAGCN only needs polynomials of the adjacency matrix with maximum degree 2 compared with 25^th^ and 12^th^ degree Laplacian matrix polynomials in Defferrard et al ^[14]^ and Levie et al ^[15]^. Though the algorithm proposing research paper evaluated the performance against spectrum filtering as well vertex domain propagation methods using three datasets, we have contrasted it using only one.

## 5. Data Integration and Filtering

### 5.1. Data Collection

The raw and unprocessed data was downloaded from multiple sources to incorporate heterogeneity of databases. The final amassed data contained drug-drug interactions, drug-protein targets and drug-disease associations. To achieve this, the databases used are:

- Drugbank ^[16]^: a reliable, continuously updated database containing information about drug targets, drug interactions, enzymes, actions etc.
- BioSNAP ^[17]^: collection of diverse biomedical networks developed by Stanford containing connections between varied biological entities from curated datasets.

The drugbank release version 5.1.8 was downloaded for availing to explore all pharmaceutical combinations that it has to offer. From BioSNAP, the networks belonging to MINER DTI dataset were considered for the study.

### 5.2. Data Preparation

We intend to utilize the drug-drug interactions and drug-protein targets provided by Drugbank. The R package “dbparser” was implemented to extract information from the drugbank XML schema. The interactions between drugs, denoted by Drugbank IDs, can easily be obtained using the dbparser::drug_interactions() method. However, the schema does not contain any direct link between said drugs and proteins denoted by standard IDs such as SwissProt or Entrez. Instead it contains descriptions of the polypeptide element, enzymes, carriers or transporters that the drugs has targeted, which is represented by a simple Biological Entity (BE) ID. Therefore, after obtaining aforementioned targets through method dbparser::targets_polypeptides(), it is necessary to map them to proteins affected by drug actions or involved in the movement of that drug across biological membranes. This results in a dataframe of our desired format, which can be utilized for the TAGCN later. Similarly, the diseases treated by drugs are derived from the Miner DG network in the Stanford Biomedical Network Dataset Collection and are denoted by MESH or OMIM IDs.

These drug-protein and drug-disease relationships are reflected as features of the drug entity during the implementation of the graph neural network, with the drug-drug interactions modelled into a graph being supplied as input.

### 5.3. Data Dimensions

The entities excerpted from the interactions between drugs produce 4294 distinctive entities. The 4294 drug entities which will become the vertices of this graph, in turn have 9631 actions with proteins and 4,16,638 associations with diseases. On examining these connections, it can be realized there are unique 2326 proteins and 5586 diseases which possess a connection with the drugs that are to be considered for the knowledge graph. This means each drug node exhibits a cumulative of 7912 features, served through the proteins and diseases. The ensuing feature vector for the neural network is an unweighted adjacency matrix with dimensions 4294×7912.

### 5.3. Graph Construction

There are 2,68,22,157 interactions between the 4294 drug nodes which serve as edges in the input graph. The graph has to be converted a simple network graph to a knowledge graph capable of holding node attributes and message embeddings using dgl, a Deep Graph Library package in Python for easy implementation of graph neural networks on top of the existing DL frameworks, here Pytorch. In accordance with this concept, the features are incorporated into the graph yielding a tensor. This also enables us to consider neighborhood topologies while predicting potential unknown interactions between existing or new drugs.

## 6. Experiments on Model

### 6.1. Topology Adaptive GCN on Knowledge Graphs

With the attempt to generalize the algorithm, the process of graph convolution has been demonstrated on the *h*-th hidden layer. Let us consider that the input feature map for each node in the graph in the *h*-th hidden layer have *F*_*h*_ features. The *f*-th feature for all the vertices in the *h*-th hidden layer are collected as input data in the form of vector 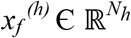, where f = 1, 2, …, *F*_*h*_ and *N*_*h*_ is the number of vertices. As present in the data graph representation, the components of *x*_*f*_ ^*(h)*^ are indexed by the vertices. Let the *g*-th graph filter be denoted by 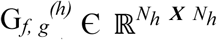. So, the graph convolution is essentially the matrix-vector product G_*f, g*_^*(h)*^*x*_*f*_ ^*(h)*^. When followed by a ReLU function, the *g*-th output feature map becomes

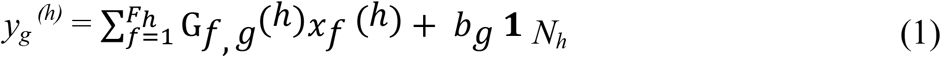

where b_g_^(h)^ is the learnable bias, and 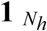 is a *N*_*h*_ dimension vector consisting of all ones. G_*f, g*_^*(h)*^ is designed such that G_*f, g*_^*(h)*^*x*_*f*_ ^*(h)*^ has a meaningful and specific convolution on a graph with a certain arbitrary topology.

As mentioned in section 4.., graph shift operation is a local operation that replaces a graph signal at a graph vertex by a linear weighted combination of the values of the graph signal at the neighboring vertices. According to this definition:

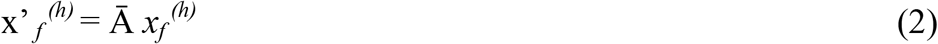

Through the graph shift Ā, along with traditional signal processing, time shift is also extended to the graph structured data. Under the appropriate assumption that G_*f, g*_^*(h)*^ is a polynomial in A, graph filter G_*f, g*_^*(h)*^ is shift invariant. To elaborate, the shift Ā and filter G_*f, g*_^*(h)*^ are commutative in the sense that Ā (G_*f, g*_^*(h)*^ *x*_*f*_ ^*(h)*^) = G_*f, g*_^*(h)*^ (Ā *x*_*f*_ ^*(h)*^).

If c _*f, g, k*_^*(h)*^ are the graph filter polynomial coefficients; the normalized adjacency matrix of the graph is quantified by A = D^-1/2^ Ā D^-1/2^, where D = diag[d] such that i-th component d(i) = ∑_*j*_*A*_*i,j*_ then it can be formulated:

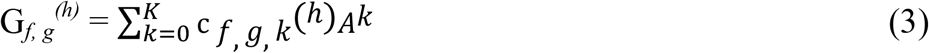

The necessity of the normalized adjacency matrix is to ensure that all eigenvalues of A are within the unit circle, and thereby G_*f, g*_^*(h)*^ is computationally stable. Next, a set of filters with different sizes are adopted (1× *F*_*h*_, 2× *F*_*h*_, …, K × *F*_*h*_)

For graph structured data, we can no longer use a square filter window as the graph topology is no longer a grid. So, 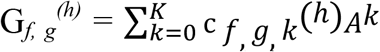 is equivalent to using a set of filters with filter size from 1 up to K. Each k sized filter, which is used for local feature extraction on the graph, is k-localized in the vertex domain.

Staying true to the standard architecture of **CNN**, a rectified linear unit is applied after every graph convolution operation as an additional non-linear function.

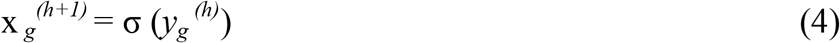

where σ (.) denotes ReLU activation function used on the graph vertices.

### 6.2. Neural Network Parameters

Among the wide range of applications that graph convolutional neural networks offer, we subscribe to the domain of link prediction. For that, it is necessary to take into consideration the types of interactions that might exist between the drugs. As discussed previously, there are seven types of existing interactions. We safely conserve 32 output features of a drug based on which the link prediction algorithm is executed. The neural network consists of 2 hidden layers with the following dimensions: (7912, 1000), (1000, 32). The other significant hyperparameter is the learning rate, which has been fixed at 0.01 after adequate tuning. In order to predict interactions between drugs, we have formulated the objective as a binary classification problem. A few requirements for efficient and informed training are to acknowledge the existence of positive and negative samples. The edges of the graph are treated as positive examples and the node pairs with no existing edges between them are considered as negative examples. These positive and negative examples are then divided into a training set and test set, followed by evaluation of the model with a binary classification metric, in this case Area Under Curve (AUC). Predictor module produces a scalar score on each edge by concatenating the incident nodes’ features and passing it to an MLP, which it does after learning the training samples through backpropagation. The separate construction of positive and negative graphs containing positive or negative examples as edges respectively makes it easier computing pairwise scores for nodes while passing messages regarding the features. Further, smooth communication of the incident nodes’ features facilitates making estimations about features of a probable edge.

After the node representation computation has been defined and the edge score computation has been attained, the overall model along with the loss function has to be established. We have resorted to a simple binary cross entropy loss calculated as:

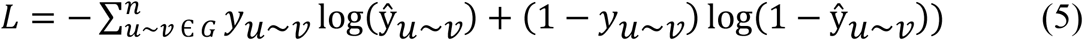

The model has been chosen as Topology Adaptive Graph Convolutional Network to integrate all the information from drugs and topological neighborhood to predict the possibility of interaction between unseen drug-drug pairs in the test set.

## 7. Results

We have evaluated the performance on the basis of Area Under Curve value which indicates the ability of the neural net model across all possible classification thresholds. The best AUC value is calculated as 97.05%. We have also observed the reduction in loss values over 500 epochs, which exhibited a steady decrease from 68.64% to 18.72% in best case scenario. The computational complexity, time and resources required for the implementation of TAGCN are also noted to be remarkably lesser than a basic GCN. The comparison between optimal performance of GCN compared to that of TAGCN is given in Table 1.

**Table 1:**
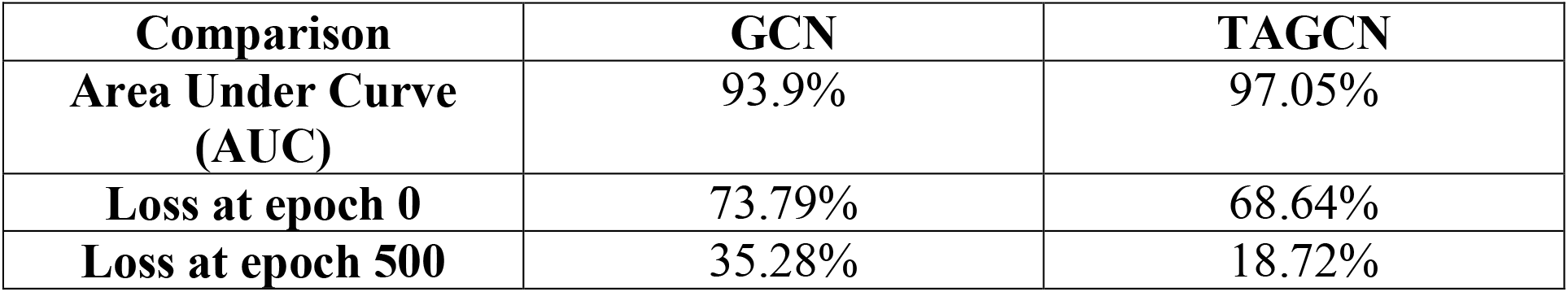
Comparative analysis between GCN and TAGCN performance.

| Comparison | GCN | TAGCN |
| --- | --- | --- |
| Area Under Curve (AUC) | 93.9% | 97.05% |
| Loss at epoch 0 | 73.79% | 68.64% |
| Loss at epoch 500 | 35.28% | 18.72% |

Figure 3 gives us the visual representation of how any particular drug is represented through all its connections in a knowledge graph context. The network centers around Bivalirudin which is shown in a rectangular hub. The triangular red nodes associated with it indicates diseases that this drug is capable of treating, whereas the yellow rectangular node indicates the protein it targets. The green elliptical nodes are few other drugs that bivalirudin is known to interact with.

**Fig 2:**
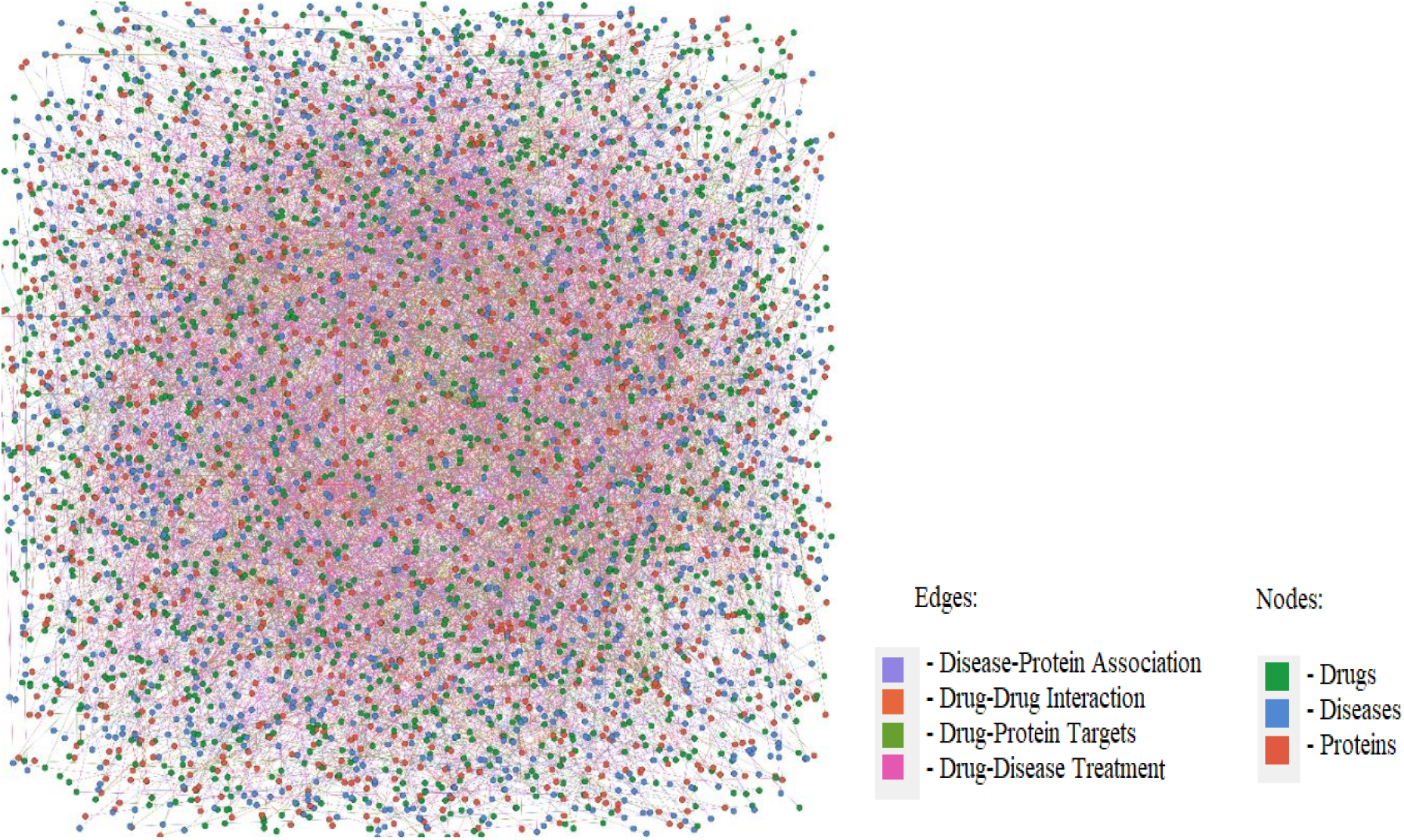
Network representation containing three types of entities and four types of interactions.

**Fig 3:**
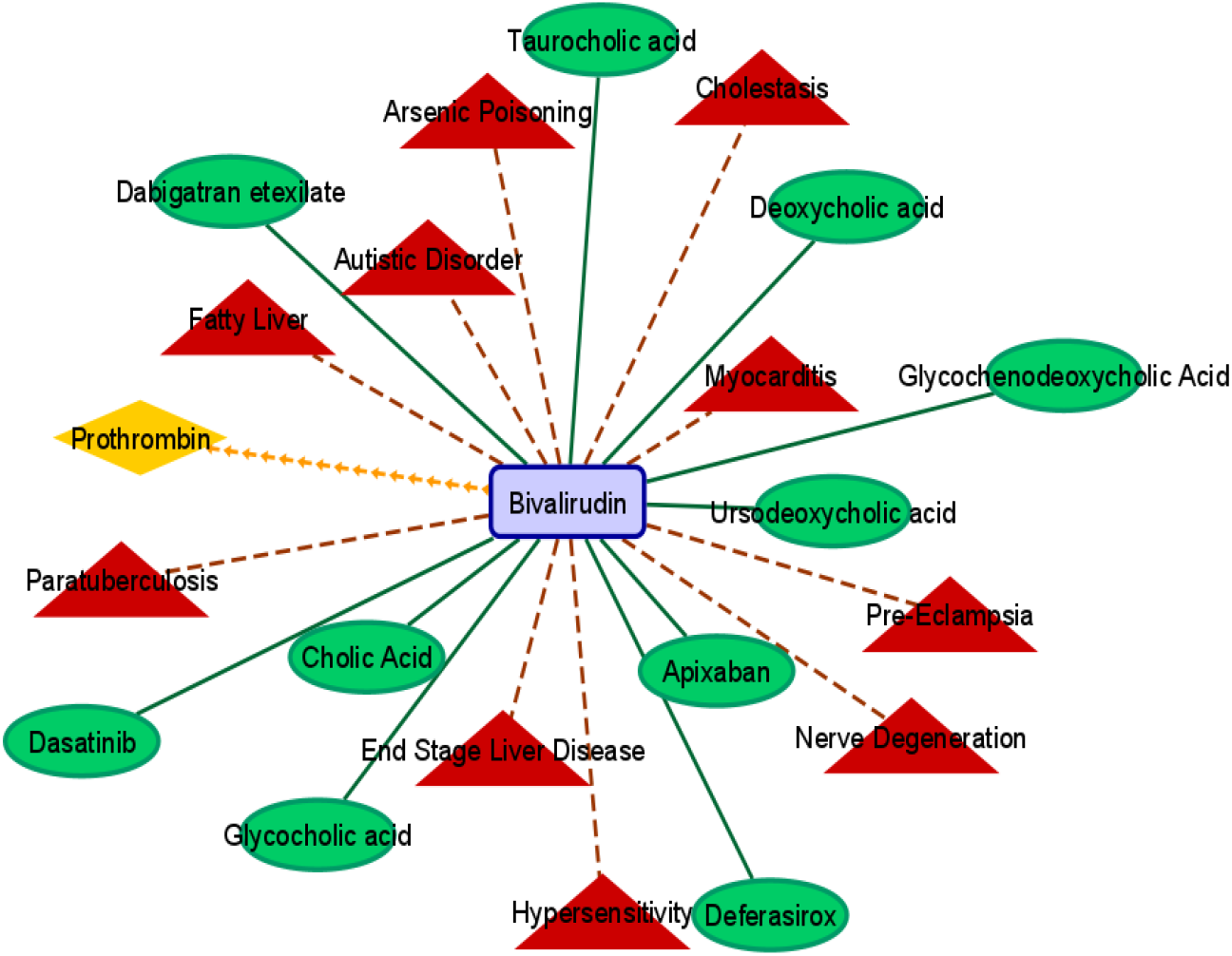
Graphical representation of the integrated entity Knowledge model of the drug ‘Bivalirudin’.

To make fair comparison, the split of training, validation and testing sets are kept in accordance with the execution by Kipf and Welling ^[18]^, i.e., labelled examples for hyperparameter tuning (filter size, dropout rate, number of hidden layers) and as mentioned before, cross entropy error is used for classification accuracy evaluation. We further compare the performance of different filter sizes from *K* = 1 to *K* = 4 in Table 2. This allows us to choose the number of hops, or rather the size of the neighbourhood we want to consider during convolution, depending on the topology of the graph. It shows that the performances for filter size *K* = 3 are always better than that for other filter sizes. The value *K* = 1 gives the worst classification accuracy. As *K* = 1 the filter is a monomial, this further validates the analysis that monomial filter results in a very rough approximation. The fact that the AUC for hop count = 4 is lesser than AUC for hop counts 2 or 3 suggests that considering a larger neighbourhood doesn’t necessarily mean better learning and prediction. It is essential to identify the optimal value for k, which translates to understanding what range interactions or receptive fields will actually enhance our learning model, before the performance flatlines or troughs.

**Table 2:** TAGCN measure of classification performance with different parameters.

| Filter Size | AUC | % Loss Reduction |
| --- | --- | --- |
| 1 | 94.76% | 67.36% down to 24.37% |
| 2 | 95.59% | 70.09% down to 22.65% |
| 3 | 97.05% | 68.6% down to 18.7% |
| 4 | 94.79% | 69.16% down to 24.84% |

In basic GCNs, the simplest convolution like operation on graphs is implemented by arranging the d-dimensional node features in an *n × d* matrix followed by node-wise transformations with feature diffusion across adjacent nodes. Here n denotes the number of nodes in the Knowledge Graph. A *L*-layer GCN model has time complexity *O(Lnd*^*2*^) and memory complexity *O (Lnd*

*+ Ld*^*2*^). However, the graph convolution in GCN is defined as a first order Chebyshev polynomial of the graph Laplacian matrix, which is an approximation to the graph convolution defined in the spectrum domain in Bruna et al ^[19]^. In contrast, graph convolution in TAGCN is rigorously defined as multiplication of polynomials of the graph adjacency matrix and not an approximation. The *K*-polynomial filters achieve a good localization in the vertex domain by integrating the node features within the *K* hop neighbourhood ^[20]^, and the number of the trainable parameters decreases to *O*(*K*) from O(n) parameters that had to learned in each layer previously. (K<<n).

## Conclusion

Drug-drug interactions are coming up as an imperative research topic because ignorance in this domain represents a considerable threat to the well-being of patients. While modifying a patient’s medical regimen, complete awareness about clinically significant interactions is advised. Graph Neural Networks come in here as connectionist models that can capture the dependence of graphs via message passing regarding side effects, chemical associations, altercations between drug nodes of the graph. Unlike standard neural networks, graph neural networks retain a state that can represent information from its neighbourhood with arbitrary depth. The application of the proposed Topology Adaptive GCN model to DDI prediction produces improved accuracy and guides us to a safer medical future.

## Future Scope

The future scopes of the thesis comprise of several directions in exploring more about the biomedical data sources. One direction is incorporating unstructured data from different sources into the proposed integrated model. The framework can be modified with more graph algorithm-based techniques. Another direction is the application of more complex deep learning architectures to understand the interaction between drug-drug-adverse events. It may include the utilization of deep neural architecture like a convolutional neural network, which can be used to extract potential predictors from a combination of different autoencoders to capture the chemical structures. The binary classification can be upgraded to a multi-label classification problem and tries to attend a comparable performance. It may also include walk/path-based graph embedding methods and 3D chemical structure as the feature of drugs. Expanding the analysis with other pairs of entities like drug-target interaction or protein-protein interactions can be studied.

Finally, aim at developing models, which is the hybridization of biomedical and machine learning techniques. Biomedical based computational models start from knowledge (in the form of theories of biochemical processes) and move towards data – a way of exploring how well current theory explains the data. On the other hand, a machine learning approach is directed at moving from data to knowledge – it outputs a mathematical model that describes the discovered relationships present within the data. Aim to design a hybrid model that combines a biomedical based model and machine learning based data-driven model to reinforce the positive aspects of both models.

## References

1. Shuman D. I., Narang S. K., Frossard P., Ortega A., Vandergheynst P. (2013). The emerging field of signal processing on graphs: Extending high-dimensional data analysis to networks and other irregular domains. IEEE Signal Processing Magazine, vol. 30, no. 3, pp. 83–98, May 2013. doi: 10.1109/MSP.2012.2235192.

2. Duvenaud D., Maclaurin D., Iparraguirre JA., Bombarelli RG., Hirzel T., Guzik AA., Adams RP. (2015). Convolutional networks on graphs for learning molecular fingerprints. In Proceedings of the 28th International Conference on Neural Information Processing Systems - Volume 2 (NIPS’15). MIT Press, Cambridge, MA, USA, Pages 2224–2232.

3. Dwivedi VP., Joshi CK., Laurent T., Bengio Y., Bresson X. (2020). Benchmarking Graph Neural Networks. 2003.00982.

4. Krompaß D., Baier S., Tresp V. (2015). Type-constrained representation learning in knowledge graphs. The Semantic Web - ISWC 2015, 9366. 1508.02593.

5. Sang S., Yang Z., Liu X., Wang L., Lin H., Wang J., Dumontier M. (2019). Gredel: A knowledge graph embedding based method for drug discovery from biomedical literatures. IEEE Access, 7:8404–8415.

6. Vilar S., Uriarte E., Santana L., Lorberbaum T., Hripcsak G., Friedman C., & Tatonetti N. P. (2014). Similarity-based modeling in large-scale prediction of drug-drug interactions. Nature protocols, 9(9), 2147–2163. doi: 10.1038/nprot.2014.151.

7. Ryu J.Y., Kim H.U., Lee S.Y. (2018). Deep learning improves prediction of drug-drug and drug-food interactions. PNAS 115(18): E4304–E4311. doi: 10.1073/pnas.1803294115.

8. Yue X., Wang Z., Huang J., Parthasarathy S., Moosavinasab S., Huang Y., Lin S.M., Zhang W., Zhang P., Sun H. (2020). Graph embedding on biomedical networks: methods, applications and evaluations. Bioinformatics, Volume 36, Issue 4, Pages 1241–1251. doi: 10.1093/bioinformatics/btz718.

9. Karim M.R., Cochez M., Chaves J.B.J., Uddin M., Oya B., Decker S. (2019). Drug-Drug Interaction Prediction Based on Knowledge Graph Embeddings and Convolutional-LSTM Network. 113–123. doi: 10.1145/3307339.3342161.

10. Zitnik M., Agrawal M., Leskovec J. (2018). Modeling polypharmacy side effects with graph convolutional networks, Bioinformatics, Volume 34, Issue 13, Pages i457–i466. doi: 10.1093/bioinformatics/bty294.

11. Asada M., Miwa M., Sasaki Y. (2018). Enhancing drug-drug interaction extraction from texts by molecular structure information. Association for Computational Linguistics, pages 680–685.

12. Du J., Zhang S., Wu G., Moura J.M.F., Kar S. (2017). Topology Adaptive Convolutional Network. 1710.10370.

13. Sandryhaila A., Moura J.M.F. (2012). Discrete signal processing on Graphs. IEEE Transactions on Signal Processing. 61(7). doi: 10.1109/TSP.2013.2238935.

14. Defferrard M., Bresson X., Vandergheynst P. (2016). Convolutional Neural Networks on Graphs with fast localized spectral filtering. Advances in Neural Information Processing Systems. 1606.09375.

15. Levie R., Monti F., Bresson X., Bronstein M.M. (2017). Cayleynets: Graph convolutional neural networks with complex rational spectral filters. 1705.07664.

16. Wishart DS., Feunang YD., Guo AC., Lo EJ, Marcu A., Grant JR., Sajed T., Johnson D., Li C., Sayeeda Z., Assempour N., Iynkkaran I., Liu Y., Maciejewski A., Gale N., Wilson A., Chin L., Cummings R., Le D., Pon .A, Knox C., Wilson M. (0217). DrugBank 5.0: a major update to the DrugBank database for 2018. Nucleic Acids Research. Nov 8. doi: 10.1093/nar/gkx1037.

17. Zitnik M., Sosi R., Maheshwari S.,Leskovec J. (2018). BioSNAP Datasets: Biomedical Network Dayaset Collection. http://snap.stanford.edu/biodata.

18. Kipf T.N., Welling M. (2017). Semi-supervised classification with graph convolutional networks. International Conference on Learning Representations.

19. Bruna J., Zaremba W., Szlam A., Lecun Y. (2013). Spectral Networks and Locally Connected Networks on Graphs.

20. Shuman D.I., Narang S.K., Frossard P., Ortega A., Vandergheynst P. The emerging field of signal processing on graphs: extending high-dimensional data analysis to networks and other irregular domains. IEEE Signal Process Mag. 2013;30(3):83–98.

